# Age-related spatial ecology of Audouin’s gull during the non-breeding season

**DOI:** 10.1101/2023.10.23.563411

**Authors:** Raquel Ponti, Virginia Morera-Pujol, Ángel Sallent, Jacob González-Solís, Raül Ramos

## Abstract

Relationships between individual’s age and the movement ecology and habitat preference of long-lived migratory birds still remain understudied. According to the exploration-refinement hypothesis it is thought that adults would select better and more productive areas for foraging than inexperienced juvenile birds would do. Additionally, age-related differences in migratory patterns and exploited habitats could be explained by the attempt to avoid competition between juveniles and adults. Here, we explored the differences in the migratory patterns, habitat selection and foraging behaviour between juvenile and adult Audouin’s gulls (*Ichthyaetus audouinii*), a species listed as vulnerable by the IUCN. We captured 9 juveniles and 8 adults in the colony of San Pedro (SE Spain) and equipped them with high-resolution 5-min programmed GPS to track their postnuptial/first migration and non-breeding destinations. First, juveniles tended to migrate longer distances than adults did. Second, the time spent foraging between age groups did not differ. Third, freshwater masses constituted an essential habitat during the non-breeding season for both juveniles and adults. Fourth, we found that adults used a greater variety of habitats than juveniles did, but adults positively select foraging habitats despite the low availability while juveniles do not. Finally, repeatability in habitat use of individuals of the same age was rather low. We provided evidence of age-related differences in migratory patterns and habitat exploitation during the non-breeding period in a migratory seabird which can be explained by the avoidance of competition between adults and juveniles and the greater experience in foraging performance that adults have in comparison with juveniles.

## Introduction

Numerous extrinsic and intrinsic factors are known to affect the spatial ecology of long-lived species. Abundant research has focused on the influence of diverse external factors on individual movement, such as environmental variables or human-induced effects [1, 2]. Conversely, little research has focused on intrinsic individual factors, such as size, age, or breeding status, as potential drivers of the movement patterns and habitat use (but see [3, 4, 5]). Possibly owing to the challenges associated with biologging juveniles, the age of the individuals is often overlooked in the literature when describing the spatial ecology of mobile species [6, 7]. In long-lived species, differences in migratory trips, foraging skills and habitat use are well noticed between juvenile and adult individuals, but less known during the non-breeding period [8, 9, 10].

Age-related differences in the spatial ecology are generally attributed to the lack of experience of juveniles, which would increase through a learning process implying a better identification of foraging sites and migratory strategies [11, 12, 13]. In long-lived, migratory species, juveniles usually perform large-scale exploratory movements until they refine their migratory route and progressively specialize their foraging sites through an “exploration-refinement” mechanism [14]. Because of the lack of experience, juveniles may perform longer daily trips during their foraging activities, increasing their foraging effort to compensate their low efficiency [15]. At the same time, juveniles tend to have high intra- and interindividual variability in movement and habitat use, which decreases as they become older and gain experience [11, 12, 13]. This ontogenetic learning process from immature stages to adulthood allows individuals to select better foraging areas and improve foraging performance, which usually entails a higher specialization of habitat use or migratory routes [16].

Alternatively, age-related differences in the spatial ecology may arise from younger individuals often migrating longer distances than adults to avoid competing directly with adults for resources and habitats [16, 17]. Age-related spatial segregation can be observed in both the breeding season, when juveniles are not hampered by breeding duties and can forage far from breeding colony, and in the non-breeding season, when juveniles and adults may shift partially or entirely their distribution to avoid intraspecific competition [12, 16, 18, 19, 20, 21].

Seabirds has been subject of extensive research with regards to their long-life expectancy and age-related variation in their foraging and movement ecology [17, 22, 23, 24]. The Audouin’s gull (*Ichthyaetus audouinii*) is a vulnerable, generalist, long-lived species, endemic of the Mediterranean Sea (Fig. 1). As a migratory gull species, it provides the opportunity to study the spatial segregation between age groups in their migratory movements and non-breeding grounds. Most Audouin’s gulls breed along the Mediterranean basin [25, 26] and non-breeding areas extend from the Western Mediterranean along the West African Coast until Senegal and Gambia [19]. Although Audouin’s gulls were traditionally considered nocturnal, specialised feeders on small pelagic fish, foraging strategies linked to rice fields and to fishery discards have often been reported [2, 27,28]. Most studies on spatial ecology of the Audouin’s gulls focus exclusively on the breeding period [2, 29, 30, 31, 32], and there is very limited knowledge on their habitat use during the non-breeding period. To our knowledge, no study has addressed age-related habitat use and foraging effort of this long-lived species, despite the interest it can have in the understanding of other seabird species’ ecology (e.g., [15, 33, 34]).

**Figure 1.**
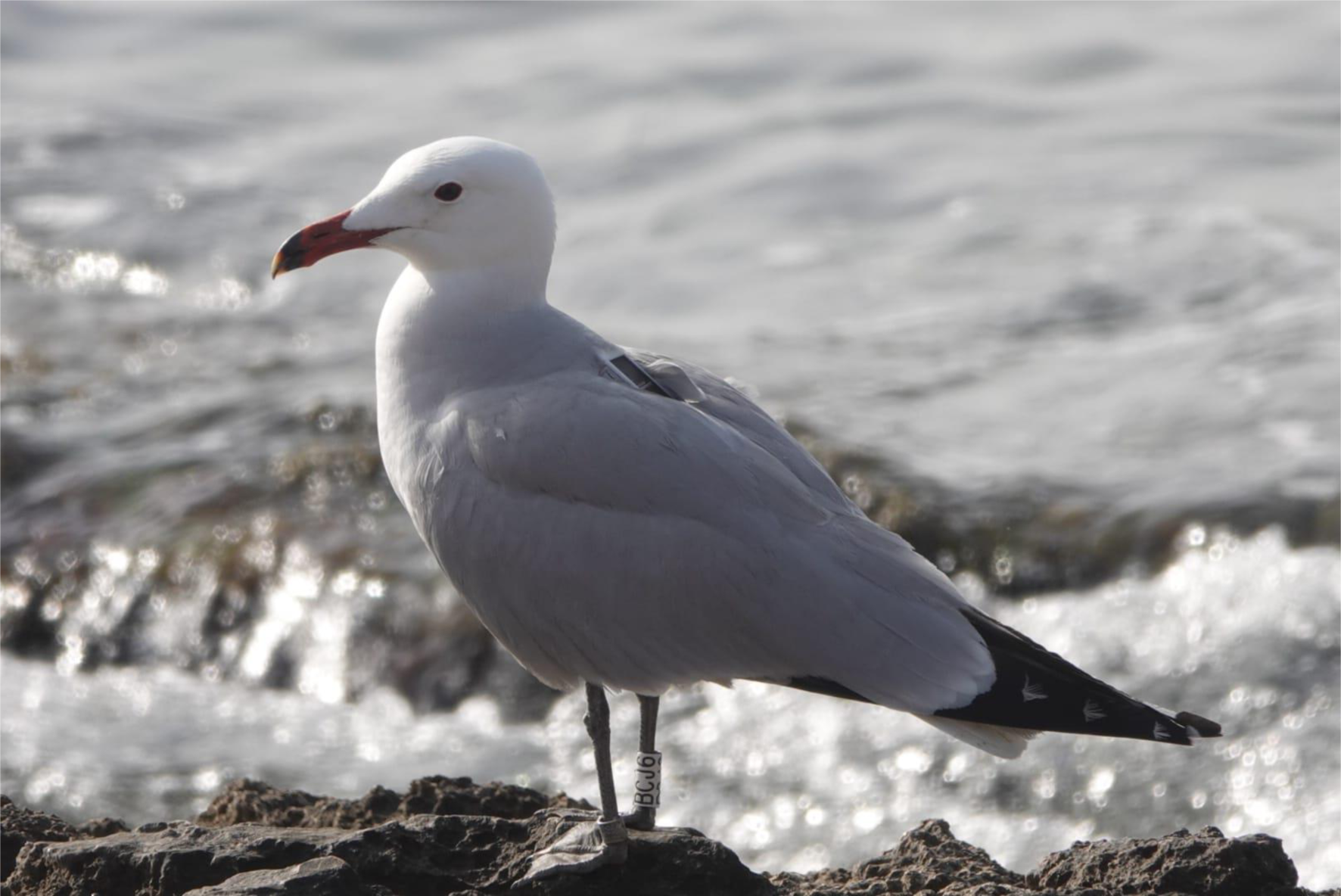
Adult of Audouin’s gull (*Ichthyaetus audouinii*) with a tracking device secured to its back with Teflon harness in Águilas (Murcia, Spain). Photo credit: Antonio Torre.

Our study mainly aims to unravel the age-related spatial ecology of the Audouin’s gull during the non-breeding season. Specifically, we are interested in determining the drivers involved in migratory distances, the flight characteristics, and the habitat selection during the non-breeding season of juvenile and adult Audouin’s gulls. We also want to explore if there were any differences in the diurnal/nocturnal activity between juveniles and adults, something that remains largely unexplored in migratory species. In accordance with the age-related segregation hypothesis [16], we expected (1) that juveniles would display a more explorative behaviour with longer migratory distances, spending more time during their foraging activities given the poorer quality of the habitat they explore. Further, and in accordance with the exploration-refinement hypothesis [14], (2) juvenile individuals would use a larger variety of habitats (i.e., larger intra-individual variability) and would be less selective in the explored habitats than adults would. Finally, (3) we also expected more inter-individual variability both in flight characteristics and habitat use in juveniles rather than in adults [16]. Thus, the juvenile explorative behaviour should derive in further differences in the type of habitat exploitation among and within juvenile individuals.

## Methods

### Fieldwork

Fieldwork was conducted in the Regional Park of *Las Salinas y Arenales de San Pedro de Pinatar* (San Pedro hereafter; latitude: 37.835°, longitude; −0.791°; Murcia, Spain) from April to June of 2020, during the incubation and chick-rearing season of Audouin’s gull. We captured 9 adults using spring traps placed on the nests and 8 juveniles by hand, just before fledgling. Birds were captured, handled and ringed with both, metal and Darvic plastic ring, under licence (*Dirección General de Medio Natural, Región de Murcia*). GPS loggers (OrniTrack 20 3G solar GPS-GSM/GPRS/3G), weighing from 17 to 20 gr, were attached on the back of birds through a harness made of Teflon ribbon.

### GPS tracking data and flight characteristic

From the 17 specimens tagged, we discarded the information of 7 (4 juveniles and 3 adults), for which the GPS information was collected during less than one month due to the failure of the device. We retained GPS information from 10 individuals, 4 juveniles and 6 adults. Audouin’s gulls start migrating from the Iberian coast towards the West African Coast (as far south as Senegal) at the end of the breeding season in June-July until the end of August [19]. Thus, we selected the months of September and October (59 days) to compare the habitat use during the non-breeding season and avoiding the migratory period.

The tracks were recorded at 1 min (70%) and 5 min resolution (30%) 24h a day and were all homogenised to the lowest resolution (i.e., 5 min) using the ‘adehabitatLT’ package for R [35], after filtering all points with invalid positions.

We calculated the home range as the 95% contour of the kernel density function using `kernelUD()` and `getvertices()` functions of ‘adehabitatHR’ package [36]. We estimated the `h` parameter through the ad hoc method `(href)` for each individual and for the age groups independently, and used the mean `h` parameter of all individuals to illustrate the individual kernels and age-grouped kernels.

We classified the tracking points per individual into three main behaviours: resting, foraging, and travelling, using the expectation-maximization binary clustering algorithm for behavioural annotation described by [37] and implemented in the ‘EMbC’ package [38]. This method differentiates four different behaviours defined by the velocity and turning angle between consecutive positions: 1) Low velocities and Low turns (LL) interpreted as *resting*, 2) Low velocities and High turns (LH) interpreted as *intensive search*, 3) High velocities and Low turns (HL) as *travelling*, and 4) High velocities and High turns (HH) as *extensive search*. In this study, we considered both, LH and HH, as *foraging* behaviour and we treated it as a unique category. The EMbC algorithm provides, for each position, the probabilities of belonging to each of the four behaviours and assigns the behaviour with the highest probability. However, we used a more conservative approach and filtered our data by retaining only the points with a minimum of 80% of probability of belonging to a certain behaviour. With this method we obtained a percentage for each behaviour per day and individual (n = 590).

Based on the tracking position obtained, we calculated for each individual the following travel characteristics: total distance (km), distance per day (km), and maximum distance to colony (km). We calculated the total distance for each individual as the sum of the distances among all tracking points (every 5 min) between September and October (n = 10), the distance per day as the sum of the distances among points during each day of tracking (n = 580), and the distance to colony as the linear distance from San Pedro to the furthest point for each individual (n = 10).

We calculated the Night Flight Index (NFI) per day and individual considering only the locations points classified as “active” behaviours (i.e., travelling and foraging), excluding the resting behaviour (n = 590). This index estimates the nocturnal/diurnal activity as the difference between the percentage of time spent in flight during the darkness and during the daylight divided by the highest value of both percentages [39]. The index ranges from −1 (only diurnal activity) to 1 (only nocturnal activity). We classified the day and the night by using the function `classify_DayTime()` of ‘RchivalTag’ package in R [40], which estimates the time period based on the timing of sunrise and sunset in each geographical area.

We compared the behavioural, flight, and NFI characteristics between juveniles and adults. We used Mann–Whitney–Wilcoxon analyses for the total distance and the maximum distance from the colony. To compare the distance per day, the NFI and the percentages of each behaviour, we performed separate Linear Mixed Models (LMM) using the age as response variable and the individual as random effect in ‘lme4’ package with the function `lmer()` in R [41], as we had several values for each individual.

### Habitat use

To evaluate habitat use we obtained from the Global Land Cover layer (years 2015 - 2019) from Copernicus Global Land Service at 100m resolution [42] the land use type at the tracking locations using the function `extract()` in ‘raster’ package for R [43]. This layer provides 23 classes of land cover, which we grouped in 7 categories: forest, herbaceous/shrubland, herbaceous wetlands, permanent water bodies, urban, agricultural areas, and ocean. We joined all forest types in one layer and the herbaceous and shrublands in another unique layer, as our study species barely occur in forests or brushy environments. We counted the habitat use for all individuals while resting, foraging, and travelling independently. We compared, within each age group (juveniles and adults), the mean percentage for all individuals of habitat use (n = 7 habitats) between behaviours (n = 3). Finally, we compared the total habitat use (considering all behaviours together) between juveniles and adults. For these calculations, we use χ^2^ using ‘MASS’ package in R [44].

### Habitat selection

We calculated, for each individual and behaviour, habitat selection by computing Manly selectivity index (MSI; [45]), which compare the habitat used to those that were available. Given that each individual exploits different areas, we used an approach where both ‘use’ and ‘availability’ are measured for each individual (III data type approach; [46]) for both juveniles and adults. First, we calculated the individual home range as the minimum convex polygon using the function `mcp()` of ‘adehabitatHR’ package [36]. Then, we used the `crop()` and `mask()` functions of ‘raster’ package [43] to obtain the Land Cover values within each individual range extent, and we randomly created 10,000 points within each individual home range using `sampleRandom()` function of ‘raster’ package [43] to represent habitat availability within the home range. From the random points obtained from the home range, we calculated the proportions for the habitats that each individual had access (i.e., habitat availability). We also transformed the number of locations per habitat and individual described in the previous section in proportions, to compare them with the habitat availability. The habitat selection was defined by the MSI (selection ratio = used/available) where values ranged from 0 to 1 indicate avoidance of the habitat studied, and values larger than 1 indicate preference or selection of the habitat. Values equal to 1 indicate non-selection neither avoidance of the habitat. We considered habitat selection when the minimum value of the 95% confidence interval is higher than 1 and habitat avoidance when the maximum value of the 95% confidence interval is lower than 1.

### Repeatability of habitat use

We calculated the Krippendorff’s alpha as estimate how repeatable individuals are in the use of habitat during the non-breeding period and in each behaviour, as has been used before in similar studies [47]. We first calculated the repeatability of habitat use between individual per age group (juveniles and adults) during each behaviour (n_juveniles_ = 4, n_adults_ = 6). This index ranges from 0, meaning individual of the same age do not select similar habitats, and 1 meaning that individuals of the same age select the same habitat. Second, we calculated the repeatability within each individual during each behaviour comparing the use of habitat per day (n = 59 per behaviour and individual). In this case, values near 0 indicate that individuals do not select the same habitats every day and values equal to 1 indicate that each individual select the same habitats during the non-breeding season. We calculated the index using the `kripp.alpha()` function of ‘irr’ package [48] in R.

### Diversity of habitat use

Finally, to test habitat use specialisation for each behaviour (foraging, resting, or travelling) at an individual and age group level, we calculated a Diversity Habitat Use index (DHU) based on Shannon index [49]: 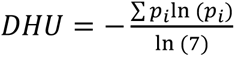, where p_i_ is the proportion of positions in each habitat. This index ranges from 0 meaning that they use only one habitat, to 1 where they equally use all the habitats.

## Results

Four out of six adults migrated to the Senegal coast, and two adults remained in the Alboran Sea (Western Mediterranean) and performed short movements within the basin (Figs 2 & S1). All juveniles migrated far from the colony they were born and remained along the African coast from the Western Sahara to Senegal. We found that, on average, juveniles performed longer trips than adults (total distance mean ± Standard Deviation [SD], juveniles: 5,922.3 ± 1,452.7 km; adults: 4,101.5 ± 699.2 km), with statistical support (n = 10, *W* = 2.0, *p*-value = 0.038). Juveniles migrated further south than adults (maximum distance from colony, juveniles: 2,909.7 ± 188.5 km; adults: 1,565.1 ± 1,031.4 km; n = 10, *W* = 0.0, *p*-value = 0.010) and travelled longer distances per day (juveniles: 98.3 ± 69.1 km; adults: 69.4 ± 46.8 km; n = 580, *t* = 2.5, *p*-value = 0.037).

**Figure 2.**
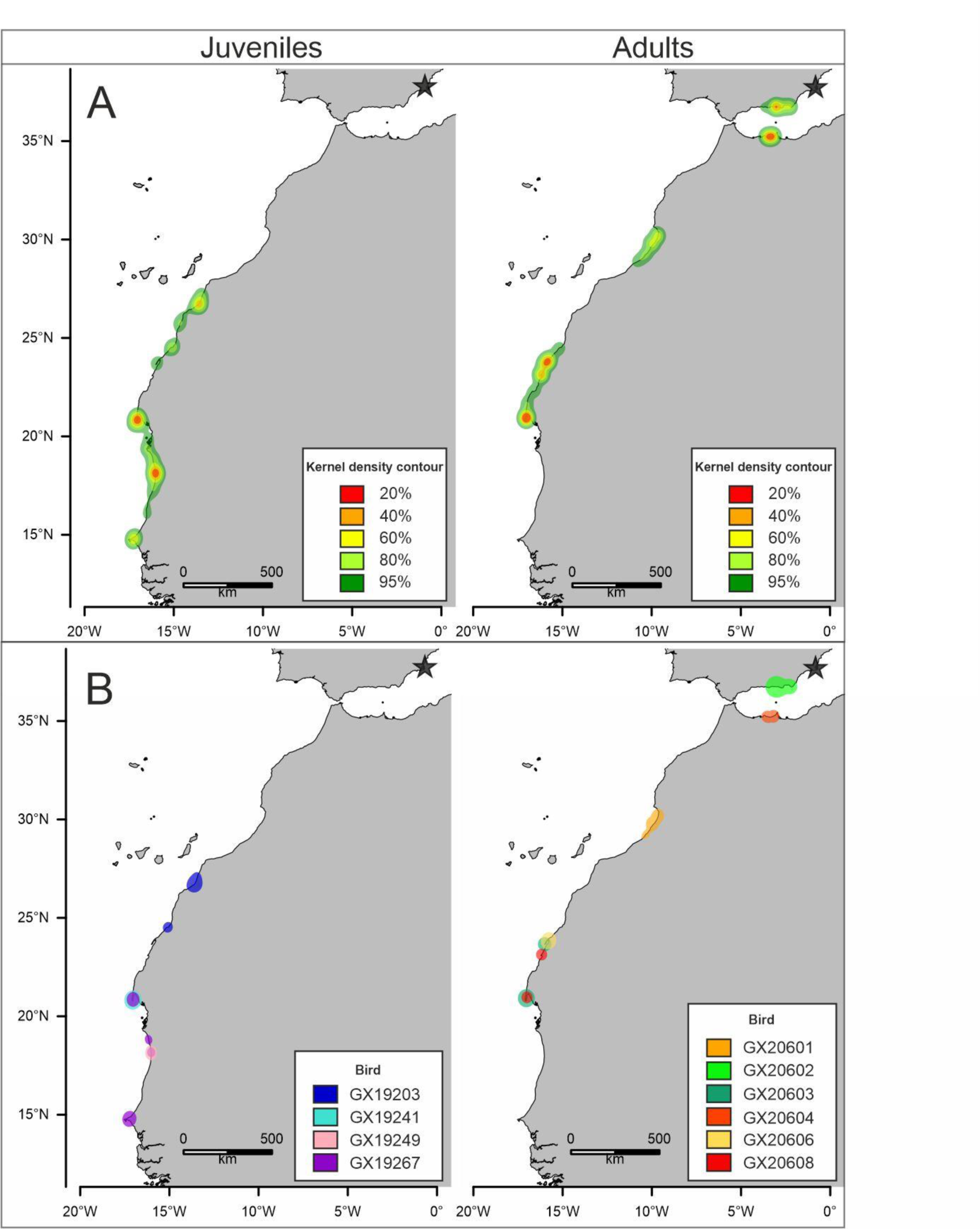
Kernel density distribution of the juveniles (A left) and adults (A right) of Audouin’s gull from September to October. 50% kernel density contours designing core areas for each juvenile (B left) and adult (B right) individual. The black star in each map indicates the location of the San Pedro colony (Spain).

Both juveniles and adults showed comparable NFI values (n = 590, *t* = −0.8 *p*-value = 0.467, Table S1, Fig. S2), indicating a slightly more diurnal behaviour for individuals in both age groups. Nevertheless, values were very close to zero, i.e. indicating an almost equal mix of diurnal and nocturnal activity (NFI: juveniles: −0.12 ± 0.03; adults: −0.07 ± 0.12).

There is no evidence of different percentage of the time spent per day in each behaviour between age groups (foraging: n = 590, *t*-value = −1.0, *p*-value = 0.335; resting: n = 590, *t*-value = 0.5, *p*-value = 0.629; travelling: n = 590, *t*-value = 1.9, *p*-value = 0.093). Both juveniles and adults spent approximately 50% of the time in foraging activities (Table 1, Table S1).

**Table 1.** Flight characteristics of juveniles and adults of Audouin’s gulls: (a) main trip characteristics: Total distance performed during September and October, distance per day, and distance to the colony, (b) Night Flight Index (NFI; from −1.0 to 1.0), (c) flight behaviour of positions (in %).

|  | Juveniles | Adults |
| --- | --- | --- |
| **Individuals** | 4 | 6 |
| | Mean ± SD | Mean ± SD |
| **a) Flight stats** |  |  |
| **Total distance (km)** | 5922.3 ± 1,452.6 | 4101.5 ± 699.2 |
| **Distance/day (km)** | 98.3 ± 69.1 | 69.4 ± 46.8 |
| **Distance to colony (km)** | 2909.7 ± 188.5 | 1565.1 ± 1031.4 |
| **b) NFI** | -0.12 ± 0.06 | -0.07 ± 0.12 |
| **c) Behavioural mode (%)** |  |  |
| **Foraging** | 49.3 ± 10.0 | 55.8 ± 9.5 |
| **Resting** | 35.8 ± 13.0 | 32.5 ± 7.7 |
| **Travelling** | 14.8 ± 3.0 | 11.8 ± 2.2 |

### Habitat use

Both, juveniles and adults, used different habitats while foraging, resting or travelling (χ^2^_juveniles_= 48.6, *p*-value < 0.001; χ^2^_adults_= 57.0, *p*-value < 0.001; Figs S3 & S4). Individuals of both ages consistently increased the use of oceanic habitats while travelling, whereas habitats such as grasslands and waterbodies were more used during foraging or resting periods (Fig. 3). The habitat use also differed among individuals of the same group in both groups, juveniles and adults (Fig. 3). Most of the differences found among individuals of the same group occurred during the foraging and resting behaviour whereas habitat use during travelling was more consistent among individuals. However, if we consider all the locations together, including all behaviours, we did not find significant differences in the habitat use between juveniles and adults (χ^2^_juvenils-adults_ = 0.1 *p*-value = 1.0).

**Figure 3.**
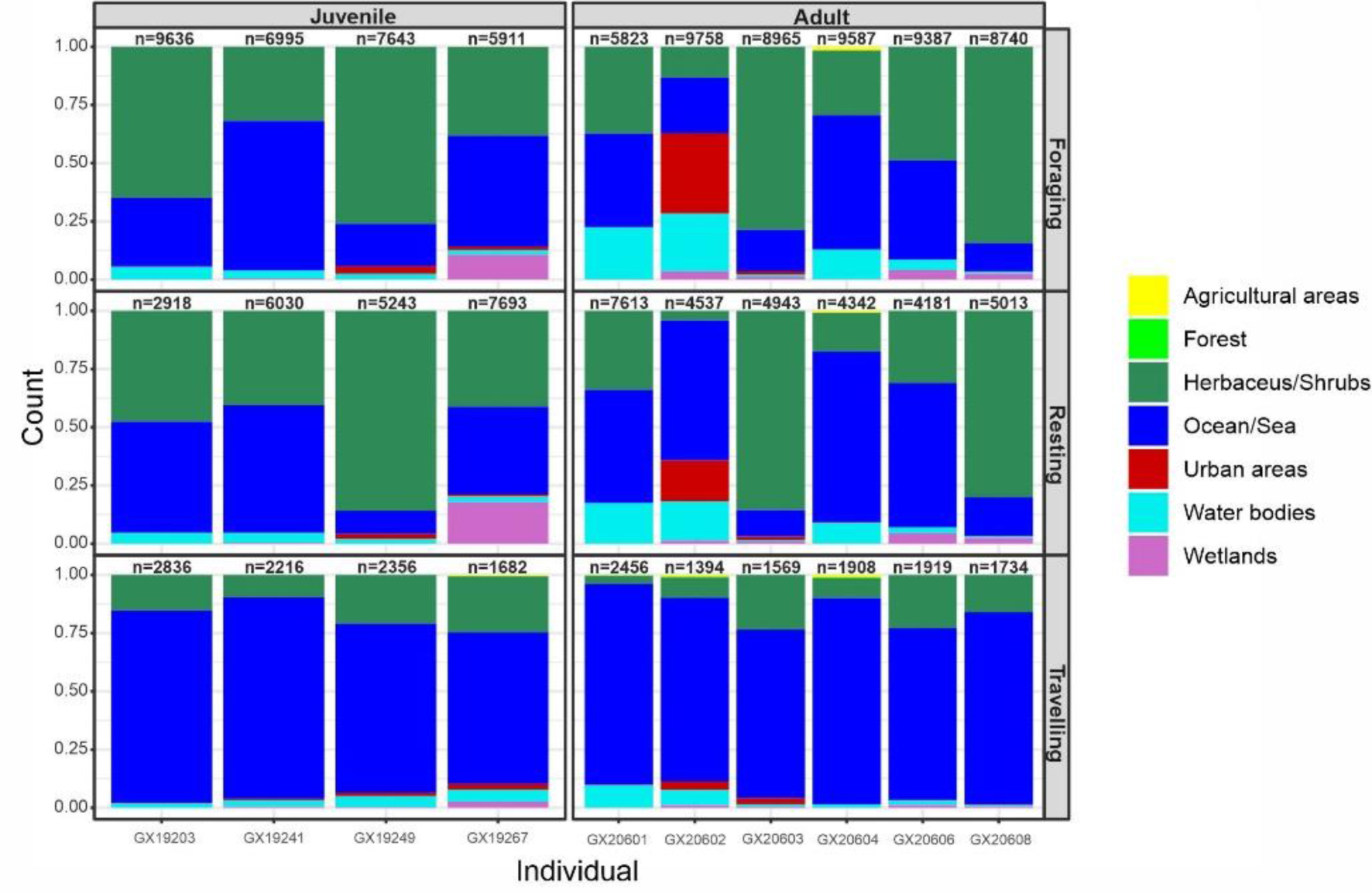
Habitat use in proportions showed by each individual and type of behaviour (foraging, resting, and travelling) for juveniles (left) and adults (right). Number of positions of each individual and type of behaviour are shown above every column (n).

### Habitat selection

MSI showed strong evidence of habitat selection (positive and negative selection) for all groups and behaviours (juveniles/adults and foraging, resting, and travelling behaviour; Table 2). During foraging activities, juveniles showed no positive selection for any habitat, such as wetlands, urban areas, grasslands, or water bodies, but negative selection for agricultural areas, forest, and oceans. However, adult gulls showed a preference for waterbodies, even though this habitat was among the less abundant habitats in their distribution. Some other habitats, such as wetlands, grasslands, or urban areas, were preferred during foraging and urban areas also during resting. Furthermore, adults refrained from selecting oceans, agricultural areas, and forests when they foraged. (Table 2). Waterbodies emerged as the most favoured habitat for both adults and juveniles. However, juveniles primarily selected it for resting, which was the sole positively favoured habitat across all behaviours, whereas adults consistently selected it in all their activities. Both juveniles and adults, avoided agricultural areas and forests in almost every behaviour. The sole exception was juveniles while travelling, as they did not exhibit a negative selection for agricultural areas. While travelling, however, gulls from both age groups tended to fly over the sea avoiding terrestrial habitats and favouring the selection of water body habitats (Table 2).

**Table 2.** Habitat selection by juvenile and adult Audouin’s gulls after the migratory period, as indicated by the MSI. Habitat selection is divided behavioural mode (foraging, resting and travelling) using **in bold**, habitats with positive selection and *in italics* habitats with negative selection. Availability: is the proportion of habitats within the home-range (50% kernel density) of juveniles or adults. Used: is habitat used by juveniles and adults. Wi: habitat selection ratio; SE: Standard Error of Wi; CI 95% confidence intervals for Wi.

|  | Juveniles FORAGING |  |  |  |  | Adults FORAGING |  |  |  |  |
| --- | --- | --- | --- | --- | --- | --- | --- | --- | --- | --- |
| Habitats | Availability | Used | Wi | SE | CI 95% | Availability | Used | Wi | SE | CI 95% |
| Herbaceous/Shrubs | 24.03 | 54.86 | 1.88 | 1.21 | [-1.07, 4.84] | **29.94** | **48.12** | **1.60** | **0.23** | **[1.03, 2.18]** |
| Agricultural areas | *0.12* | *0.02* | *0.17* | *0.11* | [-0.10, 0.43] | *3.72* | *0.28* | *0.10* | *0.15* | [-0.25, 0.46] |
| Urban areas | 0.12 | 1.13 | 11.09 | 9.60 | [-12.44, 34.62] | **0.48** | **6.69** | **13.13** | **2.90** | **[6.03, 20.23]** |
| Water bodies | 0.25 | 3.55 | 14.50 | 8.06 | [-5.25, 34.25] | **0.26** | **10.57** | **47.41** | **17.04** | **[5.65, 89.17]** |
| Wetlands | 0.18 | 2.20 | 13.02 | 14.44 | [-22.35, 48.39] | **0.04** | **2.06** | **50.48** | **19.89** | **[1.75, 99.21]** |
| Forest | *0.03* | *0.00* | *0.13* | *0.02* | [0.08, 0.18] | *0.98* | *0.02* | *0.02* | *0.03* | [-0.05, 0.09] |
| Ocean/Sea | *75.28* | *38.24* | *0.54* | *0.16* | [0.15, 0.94] | *64.58* | *32.25* | *0.49* | *0.09* | [0.28, 0.70] |
|  | Juveniles RESTING |  |  |  |  | Adults RESTING |  |  |  |  |
| Habitats | Availability | Used | Wi | SE | CI 95% | Availability | Used | Wi | SE | CI 95% |
| Herbaceous/Shrubs | 24.03 | 17.03 | 3.42 | 2.52 | [-2.74, 9.58] | 29.94 | 42.56 | 1.45 | 0.22 | [0.90, 2.00] |
| Agricultural areas | *0.12* | *0.10* | *0.00* | *0.00* | [0.00, 0.00] | *3.72* | *0.11* | *0.02* | *0.03* | [-0.06, 0.10] |
| Urban areas | 0.12 | 1.00 | 5.76 | 4.71 | [-5.79, 17.31] | **0.48** | **2.86** | **6.16** | **1.96** | **[1.36, 10.97]** |
| Water bodies | 0.25 | 3.41 | 12.97 | 6.46 | [-2.87, 28.80] | **0.26** | **8.63** | **27.18** | **7.54** | **[8.72, 45.64]** |
| Wetlands | 0.18 | 0.69 | 33.45 | 34.14 | [-50.20, 117.10] | 0.04 | 1.41 | 39.40 | 22.51 | [-15.76, 94.56] |
| Forest | *0.03* | *0.01* | *0.00* | *0.00* | [0.00, 0.00] | *0.98* | *0.01* | *0.01* | *0.01* | [-0.01, 0.02] |
| Ocean/Sea | *75.28* | *77.76* | *0.44* | *0.13* | [0.12, 0.76] | *64.58* | *44.43* | *0.70* | *0.10* | [0.45, 0.95] |
|  | Juveniles TRAVELLING |  |  |  |  | Adults TRAVELLING |  |  |  |  |
| Habitats | Availability | Used | Wi | SE | CI 95% | Availability | Used | Wi | SE | CI 95% |
| Herbaceous/Shrubs | 24.03 | 52.62 | 0.59 | 0.45 | [-0.51, 1.70] | *29.94* | *13.25* | *0.45* | *0.09* | [0.23, 0.67] |
| Agricultural areas | 0.12 | 0.00 | 1.02 | 0.69 | [-0.68, 2.71] | *3.72* | *0.35* | *0.07* | *0.07* | [-0.11, 0.25] |
| Urban areas | 0.12 | 0.89 | 10.19 | 4.14 | [0.04, 20.34] | 0.48 | 0.93 | 2.27 | 1.00 | [-0.18, 4.72] |
| Water bodies | **0.25** | **3.15** | **13.77** | **4.01** | **[3.94, 23.60]** | **0.26** | **3.82** | **13.09** | **1.74** | **[8.82, 17.36]** |
| Wetlands | 0.18 | 6.36 | 4.03 | 3.17 | [-3.73, 11.79] | 0.04 | 0.53 | 15.86 | 6.07 | [0.99, 30.73] |
| Forest | *0.03* | *0.00* | *0.44* | *0.08* | [0.26, 0.63] | *0.98* | *0.15* | *0.13* | *0.11* | [-0.14, 0.40] |
| Ocean/Sea | *75.28* | *36.99* | *1.10* | *0.34* | [0.26, 1.94] | *64.58* | *80.99* | *1.26* | *0.15* | [0.89, 1.64] |

### Repeatability of habitat use

Krippendorf’s alpha values showed that use of habitat was not repeatable among individuals (i.e., within each age group) in any behaviour. The more repeatable use of the habitat was during the resting behaviour for both, juveniles and adults (α_juveniles_ = 0.23; α_adults_ = 0.14), while foraging (α_juveniles_ = 0.07; α_adults_ = 0.11) and travelling (α_juveniles_= 0.07; α_adults_ = 0.02) showed very low repeatability.

However, overall individual repeatability values were higher when compared to those measured within age groups. During the resting period both, juveniles and adults, showed repeatable use of habitat (mean ± SD, α_juveniles_ = 0.40 ± 0.02; α_adults_= 0.38 ± 0.06). The individual Krippendorf’s alpha coefficient was also high during foraging (α_juveniles_ = 0.36 ± 0.03; α_adults_ = 0.34 ± 0.06) and travelling (α_juveniles_ = 0.30 ± 0.03; α_adults_ = 0.31 ± 0.05;).

### Diversity of habitat use

We found that Audouin’s gulls presented a high individual diversity (Table 3). Habitat use in juveniles tended to be less diverse during foraging and resting behaviours and more diverse during travelling than in adults, but differences were not significant (foraging: n = 10, W = 13.0, *p*-value = 0.914, resting: n = 10, W = 11.0, *p*-value = 0.914, travelling: n = 10, W = 10.0, *p*-value = 0.762). In case of adult gulls, the diversity of habitat use is higher during foraging, following by resting and finally travelling behaviour.

**Table 3.** (a) Individual and (b) age-related diversity index of the habitat used during foraging, resting and travelling behaviours, based on diversity Shannon indexes.

|  | Foraging | Resting | Travelling |
| --- | --- | --- | --- |
| **a) Individuals** |  |  |  |
| **GX19203 (Juvenile)** | 0.41 | 0.44 | 0.27 |
| **GX19241 (Juvenile)** | 0.40 | 0.44 | 0.26 |
| **GX19249 (Juvenile)** | 0.37 | 0.27 | 0.40 |
| **GX19267 (Juvenile)** | 0.57 | 0.60 | 0.51 |
| **GX20601 (Adult)** | 0.55 | 0.53 | 0.26 |
| **GX20602 (Adult)** | 0.74 | 0.57 | 0.41 |
| **GX20603 (Adult)** | 0.34 | 0.27 | 0.38 |
| **GX20604 (Adult)** | 0.53 | 0.40 | 0.23 |
| **GX20606 (Adult)** | 0.50 | 0.46 | 0.36 |
| **GX20608 (Adult)** | 0.27 | 0.31 | 0.27 |
| **b) Groups** |  |  |  |
| **Juveniles** | 0.49 | 0.53 | 0.36 |
| **Adults** | 0.63 | 0.57 | 0.25 |

## Discussion

The present study describes for the first time ontogenic differences in the habitat use and foraging behaviour of a gull species during the non-breeding season. We found that juveniles travel further south than adults. Also, we found that adults were more selective while foraging than juveniles, namely adults use certain habitats in more frequency than expected by the availability in their range. Although we reported a very high inter-individual variability of habitat use regardless of the age of the individuals, adults tended to exploit more diverse habitats than juveniles did, given the different habitat availability that each individual exploited.

We found that ca. 30% of the adults remained in the Mediterranean and all juveniles migrated to the West African Coast in their first migratory journey possibly to reach more productive areas and avoid competition with adults, in accordance with the age-related segregation hypothesis. As previously reported, we found that juveniles generally performed longer migrations further south than adults did. Jacob [50] and Oro and Martínez [51] noticed that most of the Audouin’s gulls that remained in the Mediterranean basin during the non-breeding period were adult individuals.

Furthermore, continuing with the age-related segregation hypothesis, we hypothesised that juveniles would spend more time foraging than adults, given their lower foraging efficiency and inexperience [15]. However, contrary to our expectations, we found no difference between juveniles and adults in the time spent foraging, resting, or travelling. While juveniles generally tend to forage less efficiently [23], they may compensate it by travelling longer distances to reach higher productive areas in southern West African Coast than adults.

The Audouin’s gull was formerly described as a nocturnal pelagic forager [27]. Our results did not reveal any diurnal or nocturnal preference for foraging at any age throughout the non-breeding season. This shows their foraging activities are not limited to the night-time and, thus, they can take advantage of fisheries or other anthropogenic activities operating during the day. Indeed, several studies have shown that daily foraging activity during the breeding period of Audouin’s gulls is mainly linked to fisheries and artificial crops like rice fields [52, 53].

Linked to the exploration – refinement hypothesis, we expected more variability of habitat use of juveniles derived by their explorative behaviour and the lack of specialization of juveniles during the first pre-breeding ages. But contrary to our first hypothesis, adults exploited a higher diversity of habitats while foraging than juveniles did, and, therefore, they would not be more specialised than juveniles would [16]. Indeed, we found that gulls showed individual preferences for some habitats regardless of their age, showing a certain degree of individual specialization within this generalist species [54]. Diversity indexes of habitat use only differed slightly between age groups and there were no differences in habitat repeatability between juveniles and adults, which suggested that juveniles and adults exploit habitats in a similar way in their non-breeding period. This pattern may be more difficult to detect during the breeding season, especially when rearing their chicks, because breeding adults behave as central place foragers, limiting their foraging ranges close to colony sites, and adults might become, therefore, more specialists [16]. Indeed, in other gull species, such as the black-backed gull (*Larus marinus*), adults exploited preferably the same habitat nearby the colony along the breeding season, and gradually tended to exploit a larger variety of habitats out of the breeding season [55]. Similarly, a high variability in habitat use can be found in populations of gulls during the non-breeding season, as observed in herring gulls (*Larus argentatus*; [56]).

Both age-groups used water bodies, defined as inland fresh or salt-water permanent bodies, as preferred habitat at least in one of the behaviours. Despite the low availability of ponds and rivers over their migratory routes and non-breeding areas, Audouin’s gulls preferentially selected this habitat. We did not find a significant positive habitat selection of juveniles during foraging. Still, it is noticed that the proportion of habitat use is higher than the availability of water bodies (Table 2). Indeed, this habitat already constitutes an important food source in breeding range of the species, possibly related to the spread of the invasive American crayfish (*Procambarus clarkia*; [57]). Since no native crayfish species is present in Africa and invasive American crayfish has not yet been detected in West Africa, this habitat would be attractive in relation to other food resources or because they represent less exposed habitats than the open ocean [53], as it was also selected for resting. No study has analysed the diet of this gull during the non-breeding season. Further studies on the topic would allow us to better understand why Audouin’s gull preferred freshwater habitats over other ecosystems. Surprisingly, both juveniles and adults avoided agricultural lands. Since rice fields are one of the most exploited habitats in the Ebro Delta in the NW Mediterranean [28, 30], we expected to find some crop preferences during the non-breeding season. However, the agricultural area classification we used included rice fields and other crops that could not be suitable for gulls which could show a non-preference of other agricultural fields.

We are aware of a few limitations of our study: the small sample size and the reduced time period could introduce some biases in our results. The low-resolution existing knowledge on land use in the African continent can also affect our findings on the preferences in habitat use. A longer time interval, including the last months of the year, would inform us about the whole non-breeding period and the temporal variability within the season. However, it is extremely challenging to gather such information given the high juvenile mortality and device limitations [19]. In addition, potential differences in the habitat use and migratory patterns could arise depending on the gulls’ colony of origin (e.g. Italy or Greece). Further investigation in this direction would provide a better understanding of whole species’ spatial ecology.

## Conclusions

Our study provides contrasting evidence for both age-related segregation hypothesis and exploration-refinement hypothesis. For instance, in relation to the age-related segregation, juveniles travelled longer distances in their migrations than adults did; but did not spend more time in foraging activities than adults did. In accordance with the explorative-refinement hypothesis, adults were more selective in their non-breeding foraging habitat than juveniles were. Conversely, our hypothesis and expectations of higher intra- and inter-variability of habitat use of juveniles in comparison to those of adults were not supported. The most likely explanation regarding the latest could be related to the difference in habitat quality of the non-breeding area of each juvenile and adult gull, which should be further explored in the future. Moreover, regardless their age, each gull showed some habitat preferences interpreted as a certain degree of individual specialization within a generalist species. Nevertheless, our approach is a basis for further investigations on the ecology of migratory gulls during the non-breeding season, which is extremely important to better understand migratory bird dynamics as an ensemble of all seasonal ecological requirements.

## Ethics approval and consent to participate

The trapping and tagging of the Audouin’s Gull in “Parque Regional de las Salinas y Arenales de San Pedro del Pinatar” were authorized by Dirección General de Medio Natural of Región de Murcia (reference: AUF/2019/0100)

## Data Availability

Raw tracking data used during the current study are available in the Seabird Tracking Database (https://data.seabirdtracking.org/) with the following code: 1757 GPS_ICHAUD_SanPedro_RRamos_2020-2021_STDB. The developed R code is freely available via the corresponding depository.

## Authors’ contributions

RP, RR and JGS conceptualized the work. AS did the fieldwork: captured the gulls and placed the GPS tracking devices. RP & VMP processed the data and performed the analyses. RP wrote the first draft of the manuscript and prepared the figures. All authors contributed to further versions of the manuscript. All authors read and approved the final manuscript.

## Acknowledgements

We would like to thank María de los Ángeles García de Alcaráz and Antonio Zamora for their help during the trapping sessions and the deployment of the GPS devices; Julio Fernández, from Salinera Española, who gave us all the facilities to work inside the salines. We also thank Antonio Torres for giving us the permit to use his picture for the Figure 1.

## List of abbreviations

DHU: Diversity Habitat Use
HH: High velocities and High turns
HL: High velocities and Low turns
LH: Low velocities and High turns
LL: Low velocities and Low turns
LMM: Linear Mixed Model
MSI: Manly Selectivity Index
NFI: Night Flight Index

