## Supplementary Material for "Age-related spatial ecology of Audouin’s gull during the non-breeding season"

Table S1. Complete Linear Mixed Models

| **Model** | N | Coefficients | Estimate (SE) | df | t-value | p-value |
| --- | --- | --- | --- | --- | --- | --- |
| Distance_per_day ~ Age + (1 \| Ring) | 580  Rings (10) | Intercept | 4.058 (0.082) | 8 | 49.5 | < 0.001 |
|  |  | Age | 0.324 (0.129) | 8 | 2.5 | 0.037 |
| NFI ~ Age + (1 \| Ring) | 590  Rings (10) | Intercept | -0.074 (0.042) | 7.9 | -1.8 | 0.113 |
|  |  | Age | -0.050 (0.066) | 7.9 | -0.8 | 0.467 |
| %Foraging ~ Age + (1 \| Ring) | 590  Rings (10) | Intercept | 0.558 (0.039) | 8 | 14.1 | < 0.001 |
|  |  | Age | -0.064 (0.063) | 8 | -1.0 | 0.335 |
| %Resting ~ Age + (1 \| Ring) | 590  Rings (10) | Intercept | 0.325 (0.041) | 8 | 7.9 | < 0.001 |
|  |  | Age | 0.033 (0.065) | 8 | 0.5 | 0.629 |
| %Travelling ~ Age + (1 \| Ring) | 590  Rings (10) | Intercept | 0.117 (0.010) | 8 | 11.1 | < 0.001 |
|  |  | Age | 0.031 (0.017) | 8 | 1.9 | 0.093 |

Figures S1. Individual tracks for all the individuals during September and October 2020.

Tracks of juvenile 1 (GX19203) and juvenile 2 (GX19241).

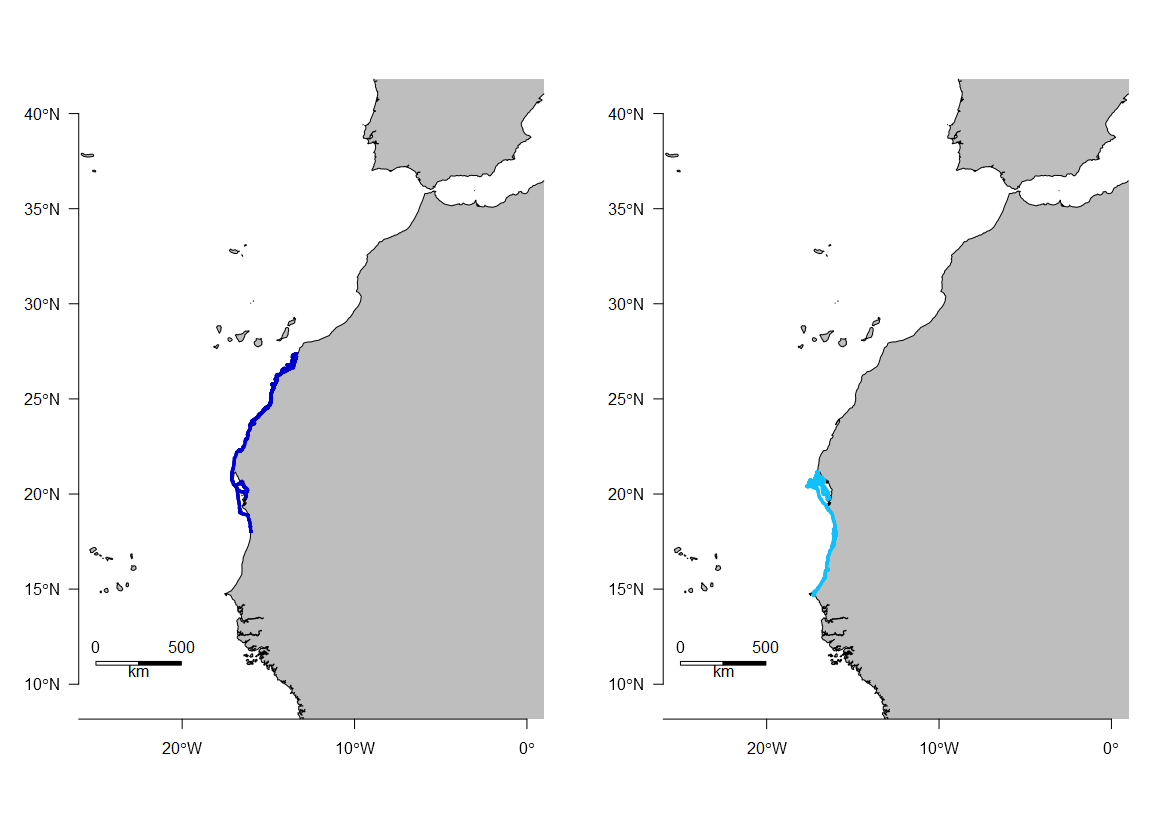

Tracks of juvenile 3 (GX19249) and juvenile 4 (GX19267).

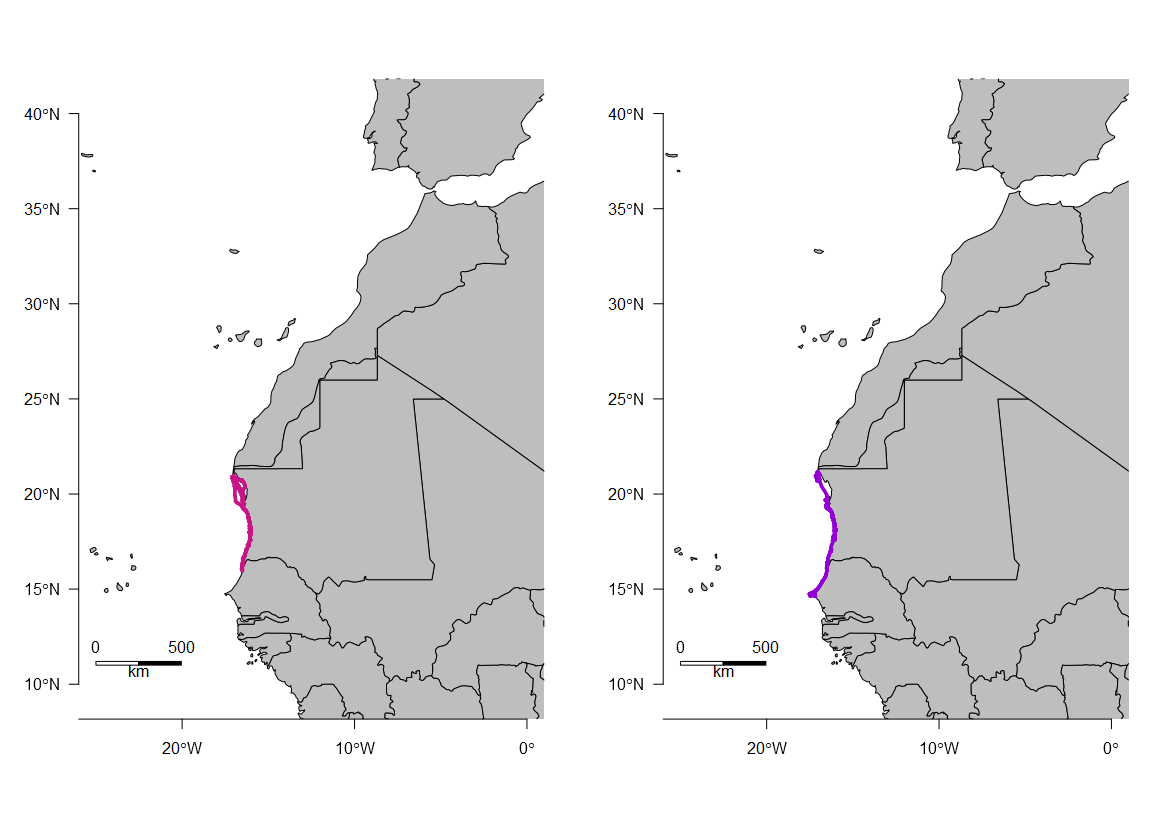
 Tracks of adult 1 (GX20601) and adult 2 (GX20602).

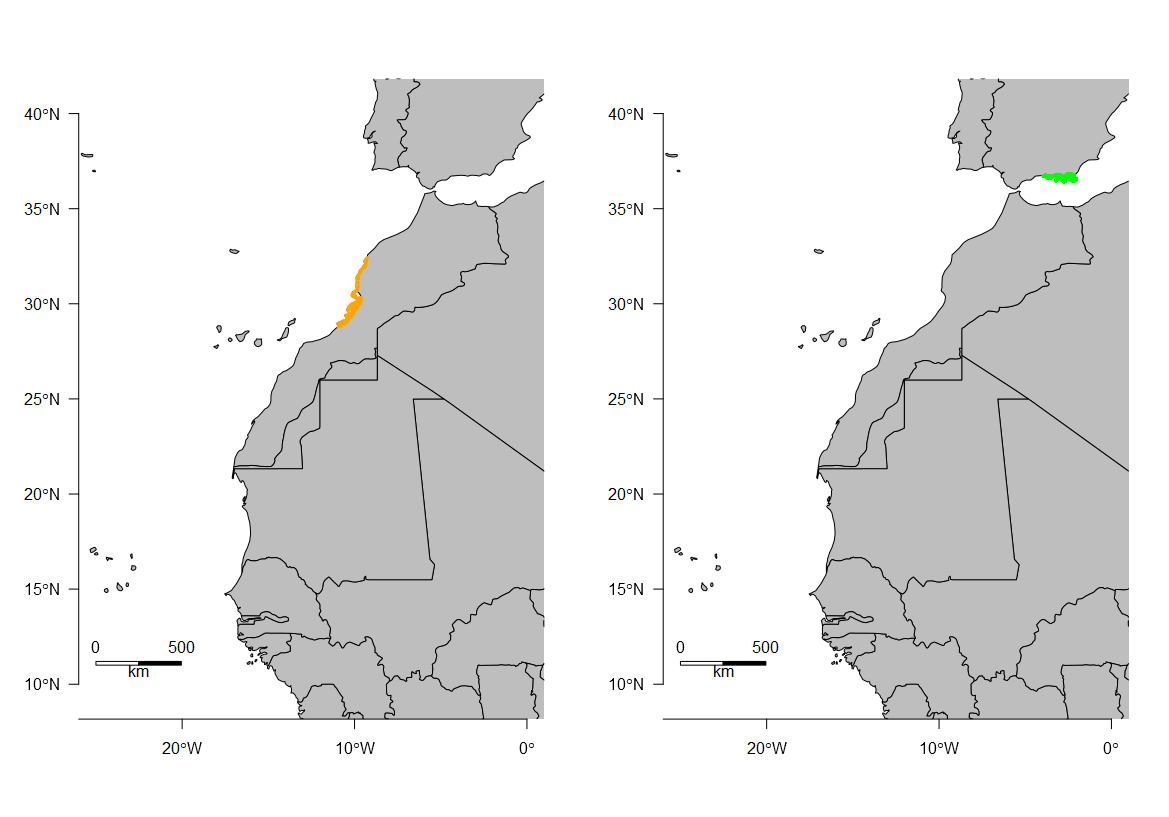

Tracks of adult 3 (GX20603) and adult 4 (GX20604).

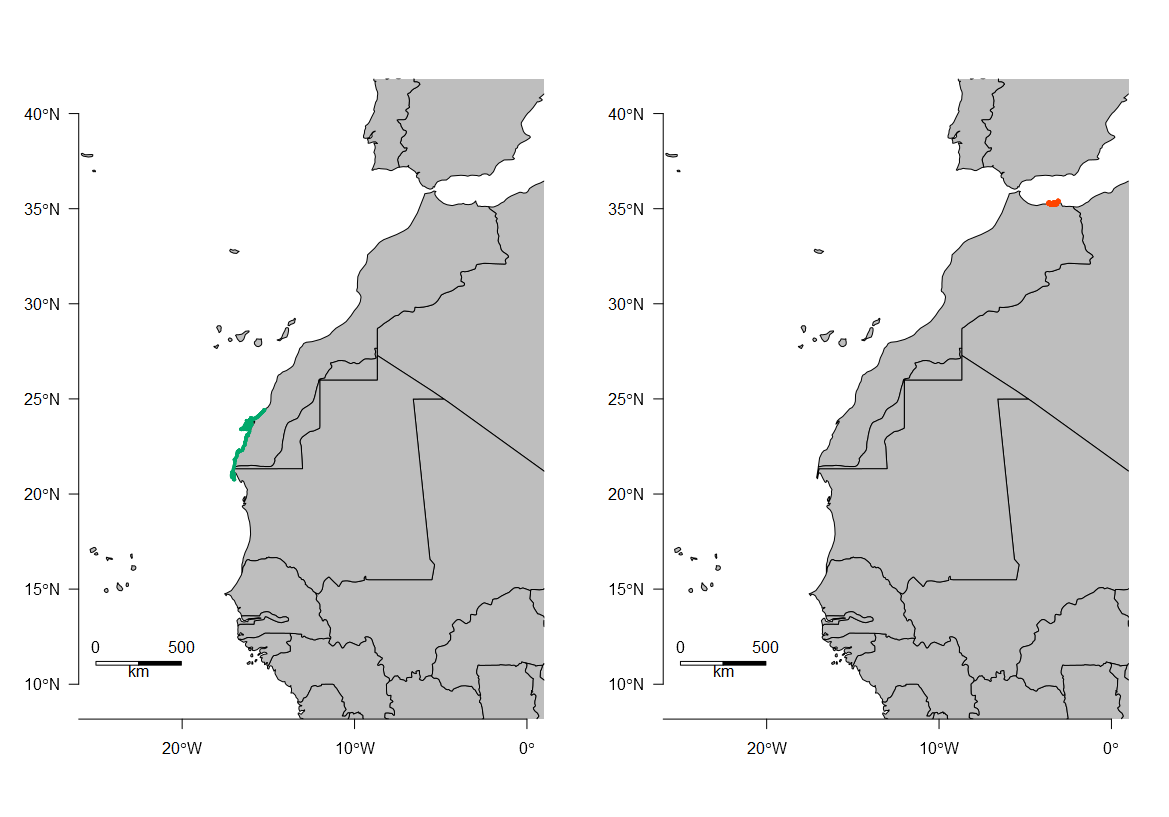
 Tracks of adult 5 (GX20606) and adult 6 (GX20608).

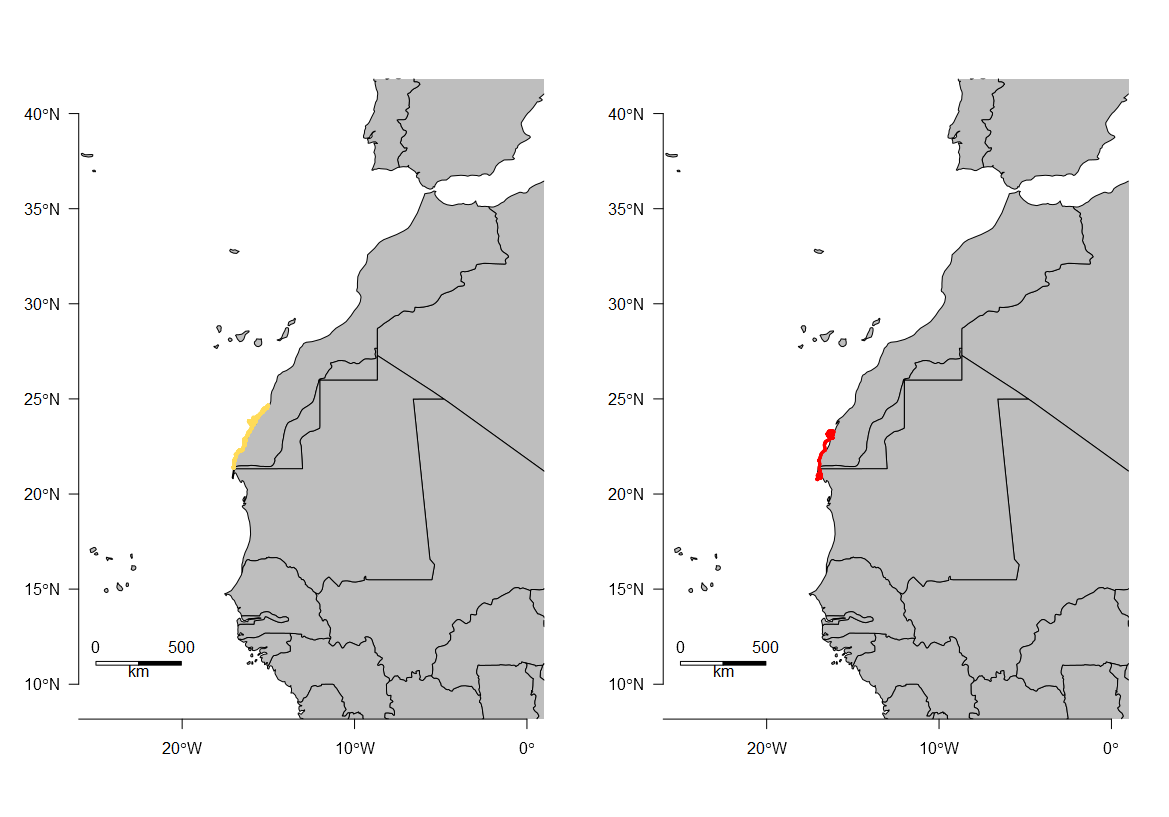

Figures S2. Density of activity as number of points with foraging and travelling behaviour per hour of the day.

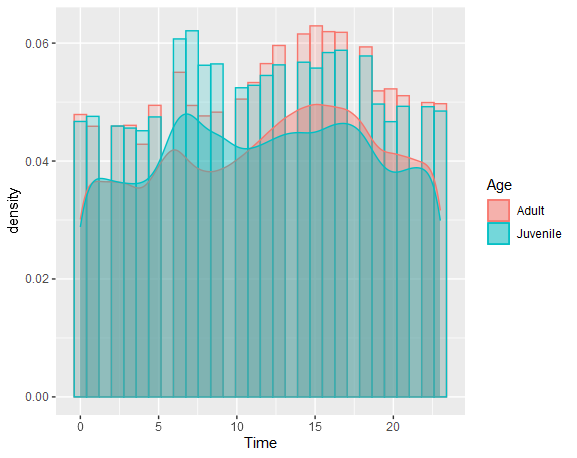

Figures S3. Habitat use per groups and behaviour, juveniles on the left and adults on the right.

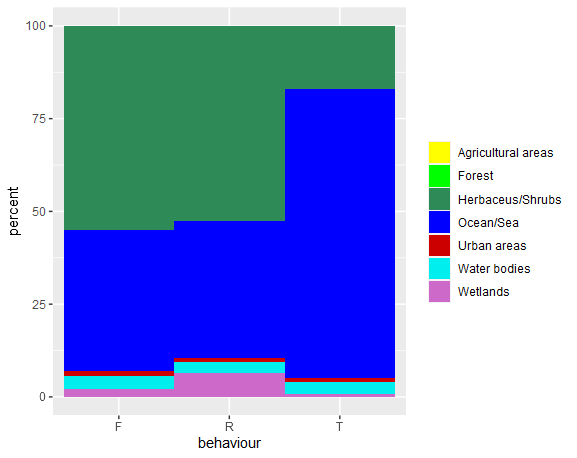

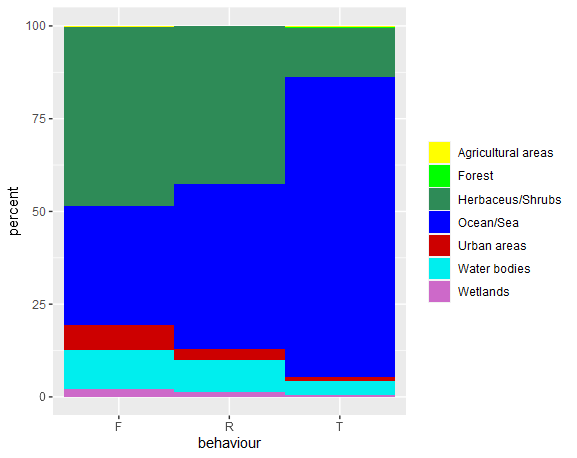

Figures S4. Tracking details of two gulls with EMbC categories defined as resident in black, foraging in golden and travelling in white. The colours of the maps correspond to the landcover categories defined in Figures S3. On the left the tracking details of a juvenile (GX19267) and on the right tracking details of an adult.

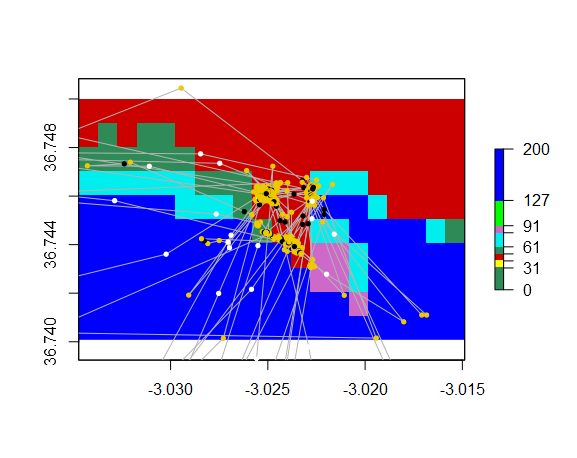

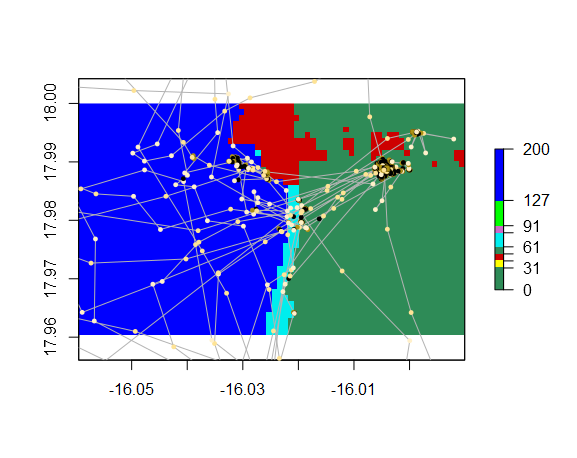
